# Investigating established EEG parameter during real-world driving

**DOI:** 10.1101/275396

**Authors:** Janna Protzak, Klaus Gramann

## Abstract

In real life, behavior is influenced by dynamically changing contextual factors and is rarely limited to simple tasks and binary choices. For a meaningful interpretation of brain dynamics underlying more natural cognitive processing in active humans, ecologically valid test scenarios are essential. To understand whether brain dynamics in restricted artificial lab settings reflect the neural activity in complex natural environments, we systematically tested the eventrelated P300 in both settings. We developed an integrative approach comprising an initial P300-study in a highly controlled laboratory set-up and a subsequent validation within a realistic driving scenario. Using a simulated dialog with a speech-based input system, increased P300 amplitudes reflected processing of infrequent and incorrect auditory feedback events in both the laboratory setting and the real world setup. Environmental noise and movement-related activity in the car driving scenario led to higher data rejection rates but revealed no effect on signal-to-noise ratio in theta and alpha frequency band or the amplitudes of the event-related P300. Our results demonstrate the possibility to investigate cognitive functions like context updating in highly adverse driving scenarios and encourage the consideration of more realistic task settings in prospective brain imaging approaches.

## 1.Introduction

To improve our understanding of human cognition and the underlying brain dynamic processes in real life situations, ecological task settings are needed that allow complex and realistic behaviors (Engel et al., 2013). On-road driving scenarios are such an example of an ecological task setting in which, in contrast to simulated driving, incorrect behavior can have drastic consequences. While laboratory studies allow controlled investigations of specific cognitive and behavioral processes, it is not clear whether these phenomena can be observed in real life conditions. This is especially the case for behaviors that involve active movements of participants which provide sensory feedback that itself influences brain dynamics and cognition (e.g. Gramann, 2013). However, the application of established brain imaging methods like electroencephalography (EEG) in more natural task settings are hindered by artefacts induced by active behavior. Non-brain activity like muscle and eye movements, or electric and mechanical artifacts can severely impact the signal quality on the sensor level. However, advances in mobile amplifier systems and developments in data analyses approaches can overcome these problems. The recently developed Mobile Brain-Body Imaging (MoBI) approach (e.g. Makeig et al., 2009; Gramann et al., 2011, 2014) overcomes the restrictions of traditional imaging modalities by using ambulatory EEG or NIRS devices combined with motion capture and other data streams that allow active behavior (e.g. Gwin et al., 2010; Jungnickel & Gramann, 2016; Banaei et al., 2017). MoBI studies demonstrate that brain activity can be distinguished from environmental and behavioral artifacts, opening up new possibilities for more realistic test and acquisition scenarios outside restricted laboratory set-ups. Driving a car is one such realistic scenario that is highly relevant for a large part of the population but represents a hostile recording environment for EEG recordings. Driving takes place in non-shielded environments with electronic equipment surrounding the driver and the task requires complex behaviors, including movement of the eyes, the head, as well as the arms and shoulders, that are typically restricted in standard laboratory settings to avoid movement-related artifacts from distorting the signal of interest. Analyzing human brain dynamics in a real driving scenario can thus be considered a stress test for comparison of EEG parameters, e.g. event-related potentials (ERP), obtained during real-world driving with parameters established in traditional laboratory settings including car simulators. If established parameters like the event-related P300 component can be replicated in real driving scenarios, EEG-data can be used to improve our understanding of how drivers process information while controlling a vehicle in a realistic environment. Providing direct access to the driver’s neuronal responses during different driving process phases, EEG might serve the development and evaluation of user centered designs for technical assistance systems in the safety-critical driving environment (e.g. Brouwer et al., 2017).

So far, only a few studies have recorded and analyzed brain activity in reallife driving tasks and the majority of these studies focus on workload measures (Kohlmorgen et al., 2007) or vigilance (e.g. Kecklund & Åkerstedt, 1993; Papadelis et al., 2007; Schmidt et al., 2009; Simon et al., 2011; Sonnleitner et al., 2014). Haufe et al. (2014) present results from a driving study for an automated braking assistance system using EEG and EMG data demonstrating the potential use of event-related potentials (ERP) to enhance automated driving technology. Because the focus of the study by Haufe and colleagues was on the replication of classification results from an earlier driving simulator study (Haufe et al., 2011), no quantitative analyses of ERP components were provided. Zhang et al. (2015) executed a combined simulator and real car study to develop a brain-computer interface (BCI) for detecting error-related EEG-activity. Despite a clear focus on classification accuracies and a small sample size for the real car experiment, the ERP results revealed comparable patterns for both acquisition scenarios, even though these were not specifically addressed in the discussion. Krol and colleagues (2017) investigated a BCI approach during interaction with an automated cruise control system in a real driving scenario. The authors demonstrate high classification accuracies for unexpected events during cruise control. However, as the focus was on classification and not replication of specific EEG features, no general conclusion can be drawn from this study about the replicability of established EEG parameters.

As no previous study has provided a detailed analysis of event related potentials during real life driving, it is still an open question whether systematic ERP-analysis is possible with data recorded in real driving scenarios and whether the results can be compared with those from traditional laboratory EEG recordings. We addressed this question by comparing the event-related P300 recorded during a dual-task driving scenario in a highly controlled laboratory setup and during an on-road driving task. The dual task scenario consisted of an interaction of the driver with a speech input device, resembling a common on-road secondary task. ERPs with onset of incorrect feedback from the speech input device were analyzed with a focus on the event-related P300 component, a positive deflection in the ERP that represents a well-established parameter for analyzing cognitive functions like attention and memory, substantiated by results from extensive laboratory assessments with numerous and heterogeneous groups of persons (Sutton et al., 1965, for reviews see Picton et al., 1992; Fabiani et al., 1987; Polich, 2007). Increased P300 amplitudes can be observed for infrequent targets in a stream of frequent stimuli (e.g. in the so-called “oddball-paradigm”). It has been argued that the reversed relationship of stimulus probability and P300 amplitudes indexes the amount of working memory updating that is necessary for the processing of the preceding stimulus (Donchin et al., 1978; Donchin & Coles, 1988) and that the P300 mediates between stimulus and response processes (Verleger et al., 2005). The P300 was expected to reflect processing of infrequent erroneous auditory feedback events in both recording environments with adequate data preprocessing in the real driving setup. Specifically, higher P300 amplitudes were expected for rare incorrect feedback events compared to correct feedback trials. In addition, the baseline EEG power spectra from both recordings were analyzed to examine possible tonic differences and to distinguish them from phasic event-related effects.

## 2. Study 1: Laboratory setup

### 2.1. Method

#### Participants

Eighteen participants volunteered for the first study. Three data sets had to be discarded due to extensive artifacts in the EEG data. The analyzed sample included 15 healthy adults (10 female, 20-35 years of age, mean 28 years). All volunteers were right handed as assessed by a German adaptation of Edinburgh handedness inventory (Oldfield, 1971) and none reported a history of neurological problems. All participants gave written informed consent and all procedures were in accordance with the principles laid out in the Declaration of Helsinki.

#### Experimental design and procedure

Participants were seated in front of a 19″ screen for visual stimulus presentation with their index fingers positioned on the marked ctrl-buttons on a standard keyboard on a table in front of them. Auditory feedback was presented through speakers placed at either side of the screen. A pool of common German first names with at least two syllables served as the stimulus material. All names were digitized as auditory feedback cues with Natural Reading Software (Natural Reading Software, Vancouver, BC Canada) and used for a simulated dialog between the driver and a technical speech based input system.

Each trial started with a black and grey flashing display for 800 ms, followed by a greyscreen for 200 ms. Three randomly chosen names from a pool of 145 forenames were presented consecutively in black letters on a grey background for 2000 ms each. In parallel, the same names were read aloud in their digitized version by a synthesized female voice. Participants were asked to remember all three names and then speak out loud the name of the numerical position that was randomly displayed at the end of the trial (e.g. “two” indicating to repeat the second name). A subsequent response interval lasted for 5000ms followed by an auditory repetition of the participant’s response. In 80 % of all cases the auditory feedback matched the stated name (eg. “Ella”), while in 20 % of all cases, only the last syllable (eg. “la”) was replayed. Correct and incorrect feedback trials were randomly presented in each trial sequence. Participants were required to wait for a tone after another 1000 ms to categorize the feedback. Correct repetitions had to be confirmed by a button press with the right index finger on the right ctrl-key and incorrect repetitions had to be indicated by pressing the left crtl-key using the left index finger (Fig. 1).

**Figure 1:**
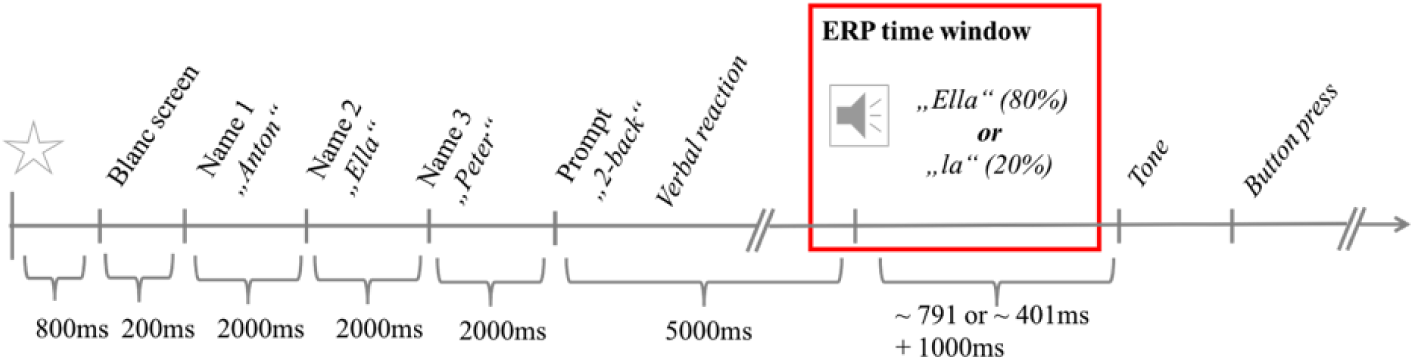
The stimulus sequence of a trial. The time interval considered for ERP-analysis is framed in red.

The task protocol followed a Wizard of Oz procedure where people believed to interact with a technical system even though operations were at least partially controlled by a human operator (cf. Dahlbäck et al., 1993). In the present case, the participants’ spoken responses were not categorized by an automated speech recognition system but by the experimental program to give a fixed error rate in the auditory feedback. Subsequent interviews revealed that none of the participants recognized the manipulation. The study consisted of six blocks of 50 trials each. The entire procedure took 2.5 h on average.

#### EEG-recording and pre-processing

EEG-data were recorded continuously from 64 active electrodes (Brainproducts GmbH, Gilching, Germany), mounted in an elastic cap according to the extended international 10-20 system (Chatrian et al., 1985), with the exception of positions PO10 and PO9, which were placed below the left and right eye respectively to measure electroocular activity. The data were digitized with a sampling rate of 1000 Hz. Prior to data recordings, impedances were brought below 5 kΩ. Off-line preprocessing and data analysis were performed in Matlab 2015 (MATLAB, The MathWorks Inc., Natick, MA, USA), using Eeglab-based routines (Delorme & Makeig, 2004). The data were filtered with a 0.1 to 100 Hz band pass filter and the sampling rate was subsequently reduced to 500Hz. Artefact contaminated channels (*M* = 11, *SD* = 3.5) were removed using automatic rejection (5 standard deviations of the mean kurtosis value or 3 standard deviations from mean probability distribution of each single channel) and subsequent manual visual inspection. Afterwards, all channels were re-referenced to an average reference calculated by the remaining channels. At this point, two copies were made of each data set. The first set was filtered with a 1 Hz high pass filter and only used for independent component analysis (ICA). The second set was filtered with a 40 Hz low pass filter and used for any further analysis. Spatially static and maximally temporally independent components (ICs) were calculated for each participant on the first set using adaptive mixture independent component analysis algorithm (AMICA, Palmer et al., 2008). The resulting ICs weighs were mapped on the 40 Hz low pass filtered sets for the ERP analysis. ICs representing eye movements were categorized for each participant (*M* = 3, *SD* = 0.6) by means of scalp maps and activation time courses. Eye movement activity was removed from the recordings by removing ocular ICs and subsequent back-projection to the sensor level.

All resulting data sets were segmented to 1800 ms epochs, starting 300 ms before the onset of the auditory feedback. For each participant, epochs were automatically discarded if amplitudes exceeded +*/-*80 *µ*V or if the measured probability of a trial exceeded a criterion of 6 standard deviations of the mean calculated probability distribution on a single channel level or 3 standard deviations for all channels. In total, 2604 correct feedback trials (*M* = 174, *SD* = 29.9) and 656 incorrect feedback trials (*M* = 44, *SD* = 6.1) were considered for the analysis.

#### Data analysis

Averaged correct and incorrect feedback amplitudes were analyzed relative to a 300 ms pre-stimulus baseline (300 – 0 ms before feedback onset). The P300 time windows and electrode sites (Fz, Cz, Pz) for analysis were selected based on the literature and visual inspection of the grand averages. A 100 ms -time window around the most positive peak at parietal electrode site Pz (738 – 838 ms after stimulus onset) was chosen for P300-analysis. Mean P300 amplitudes were assessed by 2×3 repeated measures of variance (ANOVA) with the factors feedback type (correct vs. incorrect) and electrode site (Fz, Cz, Pz). Degrees of freedom were adjusted by means of the Greenhouse-Geisser method in case of deviations from sphericicity. Post-hoc t-Tests were calculated for each condition at each electrode to evaluate differences in the topographical distribution of the measured activations and tested against correspondent Bonferroni-corrected alpha levels.

### 2.2. Results

Stimulus-locked ERP-waveforms for incorrect and correct feedback are shown in Figure 2. ANOVA results for the main P300 peak time window revealed significant main effects for feedback type, *F* (1, 14) = 59.93, *p* < .001, 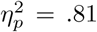 and electrode site, *F* (1.24, 17.29) = 22.11, *p* < .001, 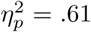. Mean P300 amplitudes were significantly higher for incorrect (*M* = 2.21, *SD* = 1.05) as compared to correct feedback (*M* = 0.07, *SD* = 0.96). Activity for both feedback conditions increased from frontal electrode site Fz (*M* = *–* 0.99, *SD* = 2.23) towards more posterior sites Cz (*M* = 1.45, *SD* = 1.23) with most prominent amplitudes at Pz (*M* = 2.97, *SD* = 0.91).

**Figure 2:**
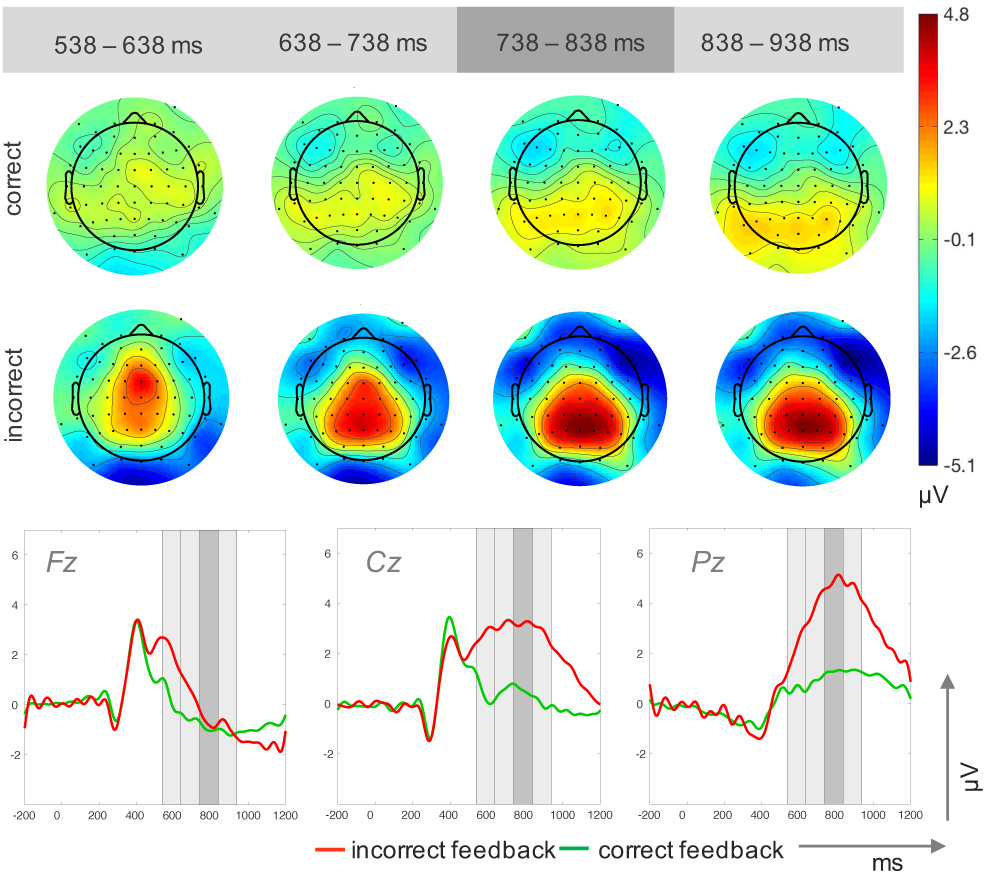
Topographic plots for four time windows (top) and ERP traces (bottom) for incorrect and correct feedback trials in Study 1. The time windows used for the upper plots are highlighted in grey in the ERP traces.

A significant electrode x feedback type interaction *F* (2, 28) = 14.65, *p* < .001, 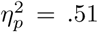 reflected amplitude differences between correct feedback trials with lower values at Fz (*M* = *–* 0.96, *SD* = 2.06) compared to Cz (*M* = 0.53, *SD* = 1.44) and compared to Pz (*M* = 1.29, *SD* = 0.62). Incorrect feedback trials resulted in reduced activity at Fz (*M* = *-*0.71, *SD* = 3.01) compared to Cz (*M* = 3.17, *SD* = 1.70) and Pz (*M* = 4.85, *SD* = 1.5). Furthermore, In correct feedback elicited larger amplitudes than correct feedback at Cz and Pz but not a frontal site Fz.

### 2.3. Discussion study 1

For the laboratory study we used a well-controlled experimental setup to establish a baseline for the experimental manipulation in the subsequent driving task. The analysis focused on the sensitivity of the P300 as an index for the processing of improbable events. As expected, the task manipulation elicited differences in event-related brain activity with increased P300 amplitudes for the infrequent incorrect feedback trials. The analysis revealed a posterior distribution with most pronounced differences between correct and incorrect feedback trials over parietal sites. This activation was absent in trials containing correct feedback information. Similarly, several studies have shown the P300 amplitude to be sensitive to stimulus probability and relevance (for a review see Polich & Herbst, 2000). In our case, the P300 appears to reflect enhanced processing costs for the categorization of the less frequent and unintended erroneous feedback events. This is in line with interpretations of the functionality of the P300 that claim that the P300 reflects the context updating within the evaluation process of new events (Donchin & Coles, 1988). In the present tasks, participants expected to hear a repeat of their own speech input. Consequently, the large P300 for incorrect fragmented feedback most likely displayed the memory update after the mismatch between the anticipated and received feedback. Moreover, as the less often incorrect feedback required a different manual button press, deviating response requirements might also be depicted by these changes (Verleger et al., 2005). The results from Study 1, confirmed our approach for investigating P300 activity for rare and deviant auditory feedback. Consequently, the procedure was applied in the following in-car recordings.

## 3. Study 2: Driving setup

Study 2 was conducted in a realistic driving setting to test whether human brain dynamics reflective of deviance detection can be recorded while participants actively drive a car. The same task as in the laboratory recordings was used to allow a direct comparison. Data processing procedures were guided by laboratory study routines reported in section 2.1.

For comparison with the data recorded in Study 1, additional data analyses were performed to answer two main questions: (1) Do changes in EEG dynamics depend on the recording environment (lab vs. car)? (2) Is there a interaction between recording environment and feedback type (incorrect vs. correct)? A main effect of feedback type should be observed irrespective of the recording environment if EEG-recordings in a driving car reliably measure brain dynamics. A main effect of recording environment would indicate an impact of the recording environment on P300 amplitudes, possibly reflecting decreased data quality due to in-vehicle artifact sources and movement of participants. Importantly, the absence of an interaction effect would indicate that the recording, analysis, and interpretation of EEG data in realistic driving scenarios is feasible for this particular task.

### 3.1. Method

#### Participants

Seventeen participants volunteered in the second study. Data from one participant had to be excluded from analysis due to technical problems during data recording, and data from a second participant had to be removed due to insufficient data quality. The analyzed sample included 15 adults (10 female, 22-36 years of age, mean 28 years,). All participants held a valid driver license for at least two years. As in Study 1, all volunteers complied with the requirements and were tested under the same conditions. None had participated in Study 1.

#### Experimental design and procedure

Participants performed the same task with identical stimulus material and time course as described in Study 1. Only set-up modifications for the incar realization are described here. The driving tests took place on a part of a restricted runway (length: approx. 1.5 *km*) of a former military airfield in Brandenburg, Germany (Fig. 3). A *Volkswagen Touran* was provided as test vehicle by the Department of Human-Machine Systems, TU Berlin. Audio feedback was transmitted through portable speakers located in the front interior. Names were presented on a 7.6 ^*"*^ TFT-display, mounted on the central console. Two buttons were added to the steering wheel, in a convenient position that allowed for safe steering and button presses with the left and right thumb. Participants were asked to maintain a speed of 40 *km/h* and the task protocol accepted deviations of +/ − 3 *km/h* (monitored via Control Area Network Data). For economic reasons and to keep up alertness, task blocks alternated with blocks in which participants worked on an acceleration and braking task, not reported here. Test blocks were defined by driving the 1.2 km test track twice back and forth (= 4.8 *km*). The total number of completed test blocks differed individually (range 12 - 14 blocks and 80 -120 trials) dependent on weather and the participant’s individual condition.

**Figure 3:**
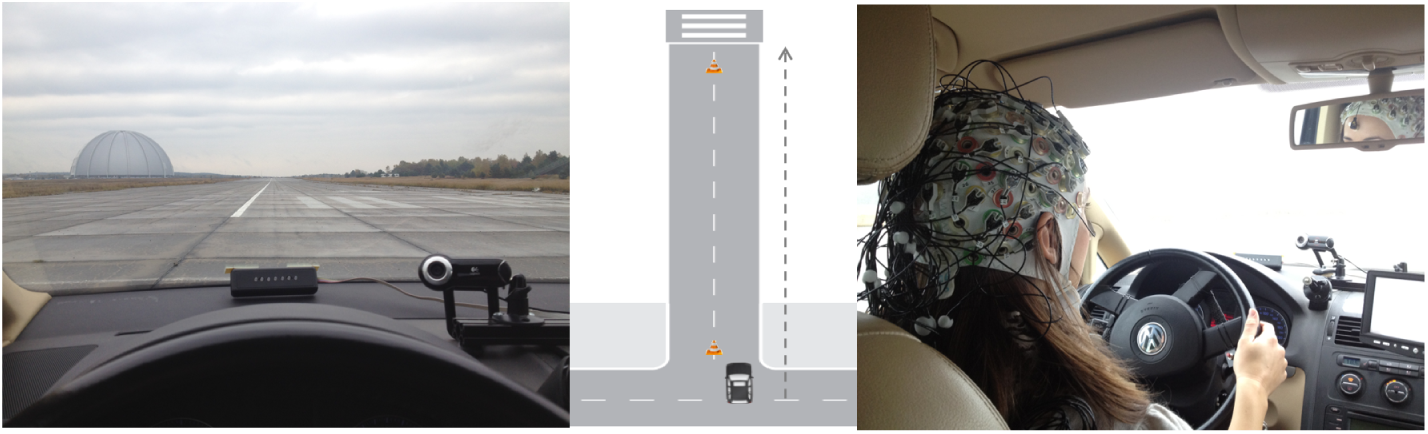
Test track from participants’ perspective (left), schematic driving course (middle) and picture from a study session (right)

#### EEG recording and preprocessing

The EEG recording setup and preprocessing steps followed the protocol for the laboratory recordings. Data were recorded with 64 active electrodes digitized with a sampling rate of 500 Hz. Impedances were kept below 5 kΩ. All data sets were offine filtered with a high-pass filter of 0.1 Hz and a low-pass filter of 40 Hz. Again, after automatic and visual inspection artifact contaminated channels were discarded (*M* = 9, *SD* = 3.1) and the remaining channels were re-referenced to an average reference. As in Study 1, two copies were made of each accordingly preprocessed data set. The first set was filtered with a 1 Hz high pass filter and only used for independent component analysis. The second data set was filtered with a 40 Hz low pass filter and used for any further reported analysis. The calculated IC weigths were map on the 40 Hz low pass filtered sets and ICs representing eye movements (*M* = 4, *SD* = 1.1) were removed. The resulting data were back projected to the channel level. Trials from the epoched data sets were automatically rejected if any channel contained amplitudes that exceeded +/ − 80 *µ*V. Slightly broader probability criterions (6 *SD* on single channel level and 3 *SD* for all channels) were applied for the automated rejection based on deviation from the mean probability distribution to adapt to the generally more fluctuating data quality of the in-car recordings. In sum, 1222 correct trials (*M* = 81, *SD* = 20.0) and 296 incorrect trials (*M* = 20, *SD* = 5.5) were considered for analysis.

#### Data analysis

Activity at midline electrodes Fz, Cz and Pz were averaged for correct and incorrect feedback trials and respectively calculated in relation to a 300 ms baseline time window preceding the auditory feedback onset. For the analysis of amplitude differences the 100 ms time widow (852 952 ms) around parietal peak activity was specified. Mean amplitudes in the P300 time-window were subjected to a 2×3 ANOVA with the factors feedback type (correct vs. incorrect) and electrode site. Greenhouse-Geisser corrections were applied and Bonferroni-corrected t-tests were calculated for post-hoc comparisons of factor levels.

Furthermore, comprehensive analysis on both data sets recorded within the two recording environments were calculated. Differences in data characteristics in terms of trial amount for both recording environments were addressed. Tonic differences in power spectrum density (*µ*V^2^*/*Hz) at midline electrode sites (Fz, Cz, Pz) were analyzed for the theta band (4 7 Hz) and alpha band (8 12 Hz). Power spectrum density estimates were calculated using Welch’s method with windows of 256 points length, zero padded to 512 points and no overlap. Mean density values were assessed for both frequency bands by a 2×3 ANOVA with factors recording environment (lab, car) and electrode site (Fz, Cz, Pz). Eventrelated amplitude differences were assessed by a 2×2×3 mixed design ANOVA with the between factor recording environment (laboratory vs. car) and the within factors feedback type (correct vs. incorrect) and electrode site.

### 3.2. Results

#### 3.2.1. Study 2

The analyses of mean amplitude values revealed a main effect for the factor feedback type, *F* (1, 14) = 31.67, *p* < .001, 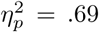, with higher amplitudes for incorrect feedback (*M* = 2.04, *SD* = 1.81) compared to correct feedback (*M* = *-*0.75, *SD* = 1.08). No further effects were found for P300 amplitudes (Fig. 4).

**Figure 4:**
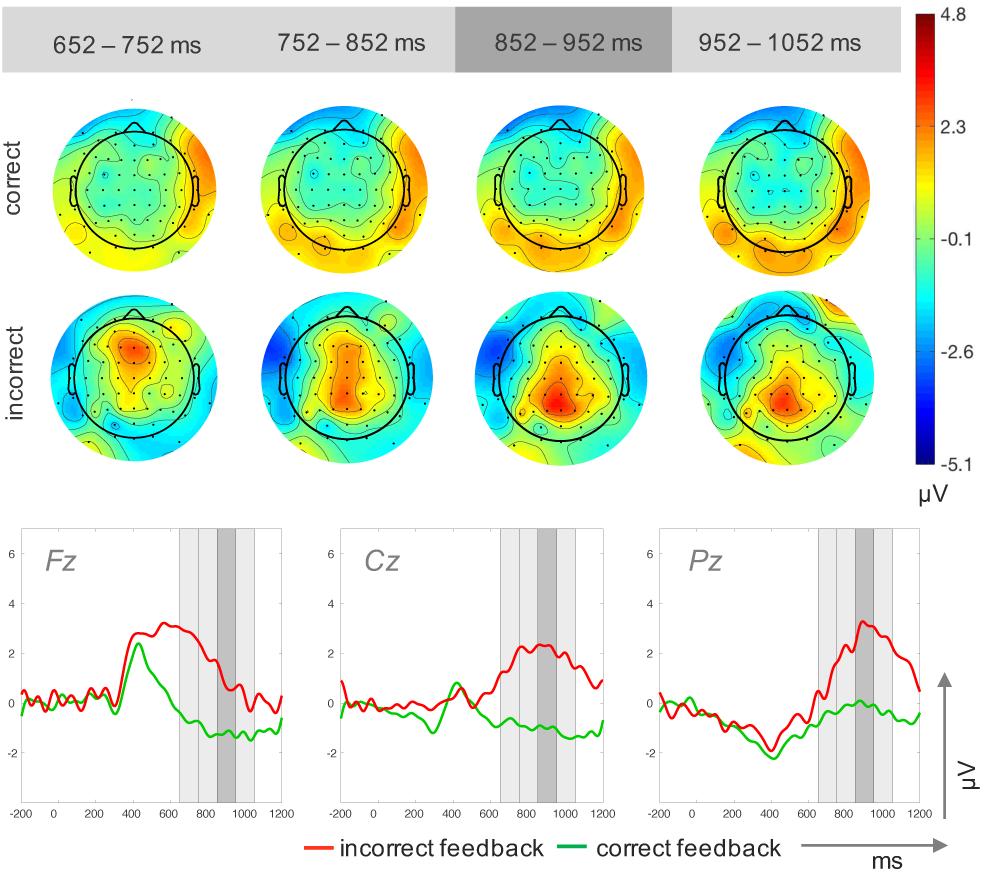
Topographic plots for four time windows (top) and ERP traces (bottom) for incorrect and correct feedback trials in Study 2. The time windows used for the upper plots are highlighted in grey in the ERP traces.

#### 3.2.2. Comparison

##### Data characteristics

In total, significantly more trials were recorded in the lab environment (3795 trials) than in the driving environment (2185 trials), *t*(21.16) = 14.11,*p <* .001. Furthermore, the proportion of trials rejected by automated cleaning was significantly higher, t(28)=-2.84, p=0.08, for epochs extracted from the driving study (28.89 %) compared to the lab recordings (14.11 %). Therefore, more trials were considered for analysis of the laboratory data (*M* = 217 trials per person, *SD* = 32.61) compared to the driving study data (*M* = 102 trials per person, *SD* = 23.30), *t*(28) = 11.05, *p* < .001.

##### Theta and alpha band power

The analysis of the power in the theta frequency band revealed a significant main effect for the factor electrode site, *F* (2, 56) = 40.76, *p* < .001, 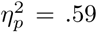 (Fig. 5). Higher theta frequencies were measured at frontal electrode site (Fz: *M* = 1.76, *SD* = 0.69) compared to the central (Cz: *M* = 0.98, *SD* = 0.44) and the parietal site (Pz: *M* = 1.02, *SD* = 0.45). With respect to the power in the alpha frequency range, a significant main effect of electrode site was observed *F* (2, 56) = 13.30, *p* < .001,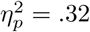 due to pronounced spectral power at frontal electrode sites (Fz: *M* = 1.02, *SD* = 0.51) and parietal electrode sites (*M* = 0.97, *SD* = 0.54) as compared to central electrode sites (*M* = 0.64, *SD* = 0.38). No main or interaction effects were found for the factor recording environment.

**Figure 5:**
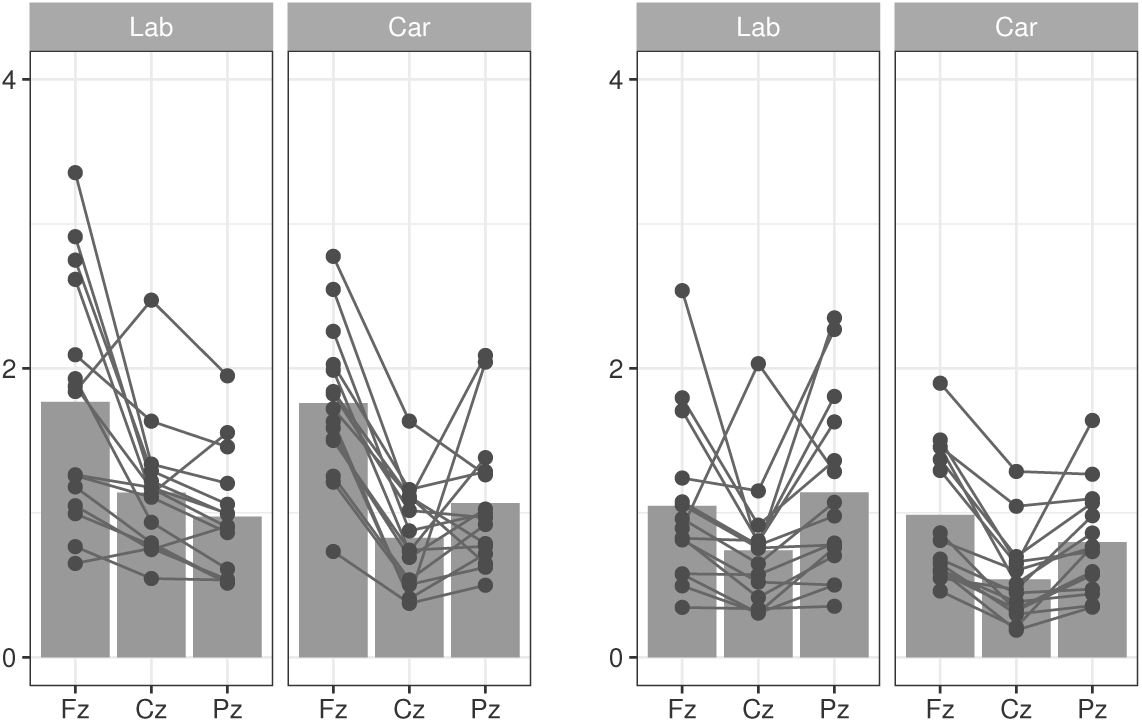
Mean Power density (y-axis, in *µ*V^2^*/*Hz) in theta (4 7 Hz, left two graphs) and alpha band (8 – 12 Hz, right two graphs) at midline electrodes (x-axis, Fz, Cz, Pz). Scatter points indicate individual mean values for each participant at each electrode.

##### ERPs

The comparison analysis on both data sets revealed significant main effects for the factor feedback type, *F* (1, 28) = 75.60, *p* < .001,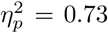, and electrode site, *F* (1.36, 38.02) = 17.40, *p* < .001, 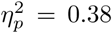. Mean P300 amplitudes elicited by incorrect feedback (*M* = 2.13, *SD* = 1.46), were more pronounced compared to correct feedback (*M* = *-*0.34, *SD* = 1.09). Activity at Fz (*M* = *-*0.59, *SD* = 2.16) was lower than activity recorded at Cz (*M* = 1.04, *SD* = 1.64) and Pz (*M* = 2.24, *SD* = 1.73).

The main effects were qualified by a significant interaction of the factors feedback type and electrode site, *F* (2, 56) = 7.84,*p* = .001, 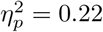, revealing highest P300 amplitudes for correct feedback at Pz (*M* = 0.62, *SD* = 1.60) compared to Fz (*M* = *-*1.09, *SD* = 2.03) but not to Cz (*M* = *-*0.22, *SD* = 1.77). Incorrect feedback elicited lower P300 amplitudes at Fz (*M* = *-*0.07, *SD* = 2.84) compared to Cz (*M* = 2.71, *SD* = 2.84) and Pz (*M* = 3.95, *SD* = 2.23). Amplitudes for incorrect feedback were larger than for correct feedback at all three electrode sites. No significant main effect or interaction effect was found for the factor recording environment.

### 3.3. Discussion Study 2

In Study 2, we tested whether the results from Study 1 could be replicated when the identical task had to be accomplished during real driving. As in the laboratory assessment, incorrect feedback elicited larger amplitudes in the P300 time window compared to correct feedback. Although ongoing parallel cognitive and motor processes are needed to solve the driving task, differences in neural response patterns for regularities and discrepancies in auditory feedback could be replicated.

In contrast to Study 1, no significant topographic variations in P300 amplitudes were found. The difference in activity patterns between the laboratory setup and the driving task might be explained by the enhanced complexity of the driving task. Frontal P300 activity for novel stimuli has been reported, to be dependent on time on task and to diminish with habituation (for a review see Friedman et al., 2001). However, less pronounced reductions in frontal activity were found for more complex tasks (Polich & Kok, 1995; Segalowitz et al., 2001). An explanation for the absence of topographic variations in the driving scenario might be due to the fact that the driving task counteracted habituation effects in the secondary task. As the driving task required constant attention, fewer resources might have been available for automated processes in the auditory secondary task.

## 4. General discussion

Two studies were conducted to establish an experimental protocol for systematically comparing the neural responses elicited by unexpected events within a realistic driving setting. In the first study we tested our experimental manipulation successfully by provoking the well-known P300 deflection for the processing of infrequent but task-relevant auditory events (Sutton et al., 1965; Katayama & Polich, 1996). In a second study, the same test was carried out in a real driving scenario, replicating the P300 response observed in the first study.

While the results demonstrate that it is feasible to investigate the neural dynamics underlying incorrect feedback processing in both scenarios, general differences in data quality had to be addressed for a more specific comparison. Despite clear visual similarities in ERP traces from both acquisitions (shown in Fig. 6), the data recorded in the car appeared to be impacted by noise. As a real-life driving scenario is an inherent source of technical and behavioral artefacts, differences in signal quality are not unexpected. This was confirmed by a significantly higher number of trials subject to automated artefact rejection due to amplitudes that exceeded a criterion of +/ − 80 *µ*V or deviated clearly from the mean calculated probability distribution. Moreover, the more complex and time consuming preparation and acquisitions sessions in the driving setup led to generally shorter recording times. These two factors accounted for a significant lower number of trials for the in-car recordings. To allow a more direct comparison, further analyses in the time and frequency domain were computed. In both studies, differences in the ERP response to correct and to incorrect feedback events were observed, while there were no differences in P300 amplitudes in incorrect feedback conditions between the two studies. The absent effect of recording environment provided the prerequisites for the comparison of both recordings. The missing interaction effect of recording environment and feedback type meant that the P300 component associated with the processing of infrequent and task-relevant stimuli was successfully replicated under realistic driving conditions. A general impact of data quality in the different recording environments on the P300 deflection can be ruled out as the tonic power spectra in both recordings were comparable and no impact of the recording environment was observed. Once more, the feasibility of EEG measurements beyond more or less restricted standard laboratory settings with new application-oriented approaches is demonstrated (e.g. Gwin et al., 2010; Debener et al., 2012; Jungnickel & Gramann, 2016).

**Figure 6:**
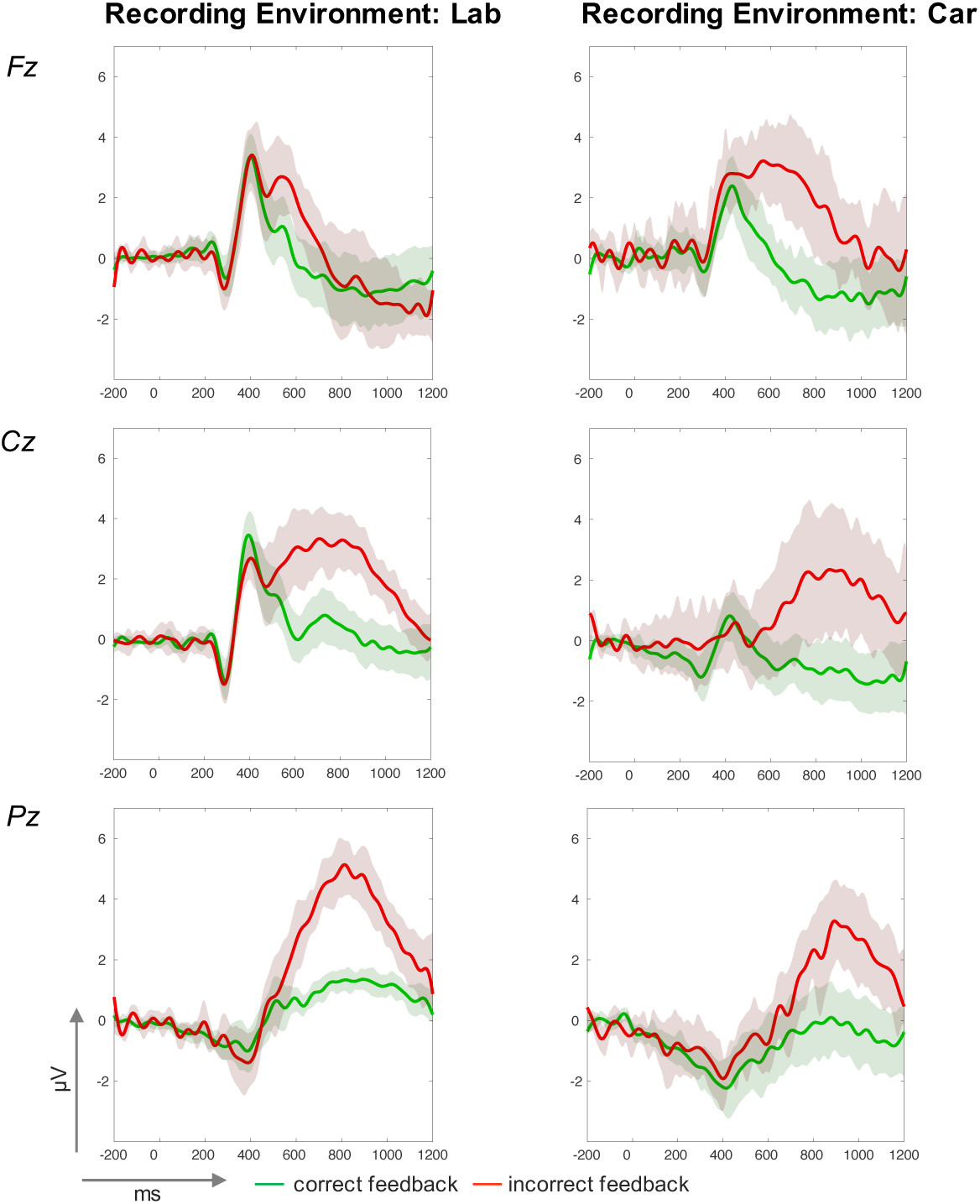
ERP-traces (in *µ*V, y-axis and time in ms, x-axis) from the laboratory (left column) and driving (right column) studies at midline electrodes Fz (top), Cz (middle), and Pz (bottom). Mean amplitude courses for correct feedback are green and for incorrect Feedback they are red. A 95%-confidence interval for each condition is indicated by the surround envelope in the corresponding color.

Despite controllable environmental influences, our approach showed furthermore that the performance of an unhindered driving task had no significant influence on P300 amplitudes elicited by unexpected and infrequent events. Based on these results, more complex dual-task paradigms with systematically varied difficulty levels in either the primary driving task or in the secondary task can be addressed. This will be of importance for further research in autonomous driving and for the development of driving assistance by providing insights into the driver’s processing of incoming information while interacting with the car and the surrounding environment. Thus, systematic analysis on variations in different stages of information processing could be used for more direct driver state assessments and the design of adaptive assistance.

## 5. Conclusion

With two studies we were first able to replicate previous laboratory- based work on P300 amplitudes and then to confirm a high level of ecological validity of our results in a realistic driving task setting. Our findings provide strong evidence that complex cognitive functions like context and response updating processes can be examined in a highly adverse driving environment. The processing of infrequent and incorrect auditory feedback events was reflected by comparable P300 patterns in both recordings. More specifically, amplitudes and tonic EEG power spectra from both studies were not affected by the recording environment. The possibilities to provide direct insights into brain dynamics of humans participating in a real world driving task provides compelling arguments for further investigation in realistic task settings with more complex manipulation or on less robust potentials. A gradual transfer of the extensive knowledge gathered from laboratory ERP reports into ecological task settings could prospectively result in complex findings about brain dynamics of actively behaving humans.

